# IBDome: An integrated molecular, histopathological, and clinical atlas of inflammatory bowel diseases

**DOI:** 10.1101/2025.03.26.645544

**Authors:** Christina Plattner, Gregor Sturm, Anja A. Kühl, Raja Atreya, Sandro Carollo, Raphael Gronauer, Dietmar Rieder, Michael Günther, Steffen Ormanns, Claudia Manzl, Stefan Wirtz, Asier Rabasco Meneghetti, Ahmed N. Hegazy, Jay V. Patankar, Zunamys I. Carrero, TRR241 IBDome Consortium, Markus F. Neurath, Jakob Nikolas Kather, Christoph Becker, Britta Siegmund, Zlatko Trajanoski

## Abstract

Multi-omic and multimodal datasets with detailed clinical annotations offer significant potential to advance our understanding of inflammatory bowel diseases (IBD), refine diagnostics, and enable personalized therapeutic strategies. In this multi-cohort study, we performed an extensive multi-omic and multimodal analysis of 1,002 clinically annotated patients with IBD and non-IBD controls, incorporating whole-exome and RNA sequencing of normal and inflamed gut tissues, serum proteomics, and histopathological assessments from images of H&E-stained tissue sections. Transcriptomic profiles of normal and inflamed tissues revealed distinct site-specific inflammatory signatures in Crohn’s disease (CD) and ulcerative colitis (UC). Leveraging serum proteomics, we developed an inflammatory protein severity signature that reflects underlying intestinal molecular inflammation. Furthermore, foundation model-based deep learning accurately predicted histologic disease activity scores from images of H&E-stained intestinal tissue sections, offering a robust tool for clinical evaluation. Our integrative analysis highlights the potential of combining multi-omics and advanced computational approaches to improve our understanding and management of IBD.

## Introduction

Inflammatory bowel disease (IBD) is a non-infectious chronic inflammatory disease of the gastrointestinal (GI) tract. It manifests as two major subtypes, ulcerative colitis (UC) and Crohn’s disease (CD). In UC, the inflammation is limited to the mucosa and submucosa of the colon and continuously spreads to a varying extent from the rectum to the proximal colon and, in severe cases, to the terminal ileum (backwash ileitis). CD affects all layers of the gastrointestinal wall and may discontinuously affect different portions of the entire GI tract. Symptoms of both UC and CD include diarrhea, rectal bleeding, abdominal pain, weight loss, and fatigue^1^. IBD increases the risk of colorectal cancer^2^, and of concomitant manifestation of other immune-mediated inflammatory conditions, such as arthritis^3^. The disease affected 4.9 million persons worldwide in 2019, and both incidence and prevalence have been increasing globally since 1990^4^. The exact cause of the disease is currently not known, but the leading hypothesis is that it arises from a combination of genetic predisposition, dysbiosis of the gut microbiome, and environmental factors, that lead to excessive activation of the mucosal immune system^5^.

Despite recent advances in the treatment of IBD patients, including the development of advanced targeted therapies, IBD can currently not be cured. Therefore, clinical interventions focus on minimizing symptoms with immunosuppressive and anti-inflammatory drugs^1^. First-line treatment options include aminosalicylates for mild cases of UC and various steroid prparations for mild to severe cases of UC and CD. More recently, targeted therapies such as tumor necrosis factor α (TNFα) inhibitors are being used in moderate to severe cases, with promising results^1,6^. However, about a third of IBD patients are refractory to anti-TNFα treatment, and of the primary responders, 23-46% lose their response per year^6^. Patients failing to respond to treatment may require surgical removal of the inflamed intestinal segments^1^. Clinical symptoms do not always reliably reflect disease activity, as patients may experience significant inflammation without overt symptoms or report severe symptoms despite minimal inflammatory activity. This inconsistency underscores the need for objective measures of disease activity to guide clinical decision-making and improve patient outcomes. However, there is no single “gold standard” for diagnosing IBD, assessing disease severity, or evaluating treatment response. A multifaceted approach is employed by physicians, integrating clinical symptoms, laboratory biomarkers, radiological imaging, endoscopic examinations, and histological analysis of biopsy specimens^7^. While this comprehensive strategy provides valuable insights, it also highlights the complexities of assessing disease activity and the ongoing need for standardized, objective, and accessible diagnostic tools.

In this study, we address these challenges by creating a comprehensive multi-center, multi-omic, and multimodal IBD atlas (IBDome atlas), integrating individual genomic, transcriptomic, proteomic, histopathologic, and clinical data from 1,002 IBD patients and respective controls. Using this resource, we investigate site-specific immunological pathways and features, develop a novel serum protein-based disease activity signature (IBD-IPSS), and leverage deep learning prediction of histologic disease activity from histological images through the use of general-purpose foundation models. Our integrative approach aims to provide a more comprehensive understanding of the IBD immunopathogenesis, by combining detailed clinical disease characteristics and in-depth multi-omic molecular analyses on an individual level in a multi-modal IBD atlas, enabling novel translational research approaches and pathophysiological concepts that will foster the concept of personalized medicine in IBD.

## Results

### Development of the IBDome atlas

We first generated multi-omic and multimodal data, encompassing clinical metadata from 1,002 patients diagnosed with IBD and a matched cohort of individuals without IBD including histopathology, high-resolution H&E images, whole exome sequencing (WES), RNA-sequencing, serum proteomics data, endoscopic activity scores, stool appearance scores, and clinical disease characteristics to comprehensively characterize the underlying immunopathogenesis of IBD in the individual patient (**Fig. 1a, b and Extended Data Fig. 1a, b**). We consolidated all datasets into a unified relational database, termed the IBDome atlas. In total, this atlas includes data from 539 patients diagnosed with CD, 321 patients with UC, 26 patients with indeterminate colitis (IC), and 116 non-IBD controls without any intestinal inflammatory condition from two distinct study centers, Berlin and Erlangen (**Fig. 1c**). To facilitate the exploration of the clinical and molecular data, we developed an interactive and publicly available web application, accessible at https://ibdome.org. The graphical user interface allows to interactively select patients based on clinical variables and visualize gene expression or correlation with protein abundance, endoscopy and histopathology scores.

**Fig. 1.**
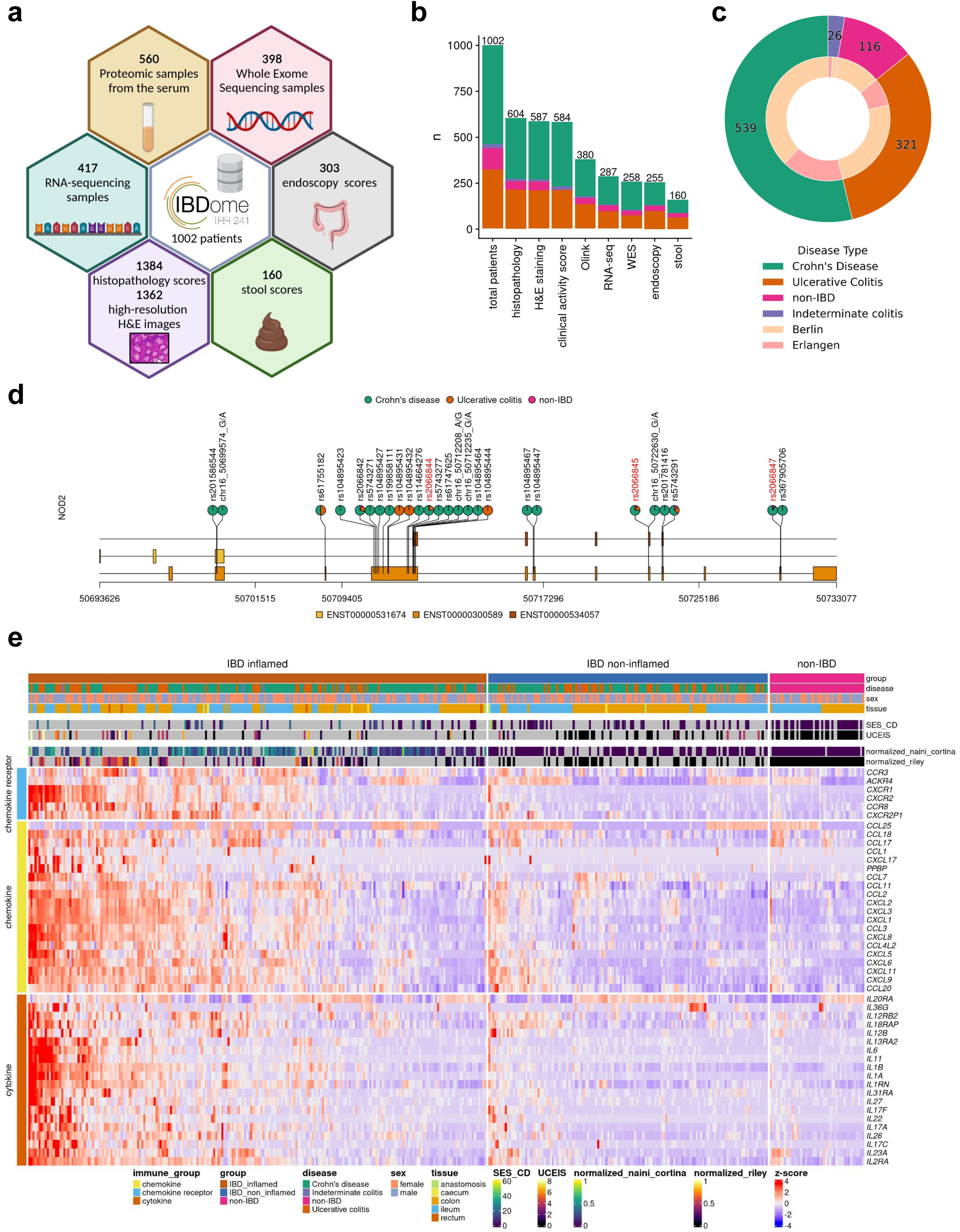
Characteristics of the IBDome atlas. **a,** Schematic overview of the datasets and sample numbers for the 1002 patients integrated in IBDome. **b,** Number of patients per sample type; colors are representing the different diseases and numbers on top of the graphs are depicting the total numbers. **c,** Patient distribution illustrated as a nested pie chart, with the outer circle representing the number of patients per disease and the inner circle indicating the proportion of patients per study center (Berlin and Erlangen). **d,** Exome mutation map of NOD2; highlighted in red are the known most frequent variants R702W (rs2066844), G908R (rs2066845), and 1007fs (rs2066847). **e,** Heatmap of differentially expressed cytokines, chemokines, and chemokine receptors between IBD inflamed samples (n=223) versus non-IBD controls (n=46), clustered by euclidean distance and complete linkage. SES-CD = Simple Endoscopic Score for Crohn’s Disease; UCEIS = Ulcerative Colitis Endoscopic Index of Severity.

Genomic and transcriptomic characterization confirms that our atlas accurately represents the molecular landscape of IBD (**Fig. 1d, e**). As expected, mutations in *NOD2* are predominantly observed in CD patients. The three most common variants (R702W, G908R, and 1007fs)^8^ exhibit higher mutation frequencies compared to UC and non-IBD patients (**Fig. 1d**). Differential expression analysis between inflamed IBD (tissue- and date-matching histopathology score > 0) and non-IBD control samples showed an upregulation of cytokines, chemokines, and chemokine receptors associated with disease severity scores determined by histopathology or endoscopy scores (**Fig. 1e, Extended Data Table 1**). Furthermore, disease activity scores (modified Naini Cortina^9^ and modified Riley^10^ scores evaluated through histopathology, UCEIS - Ulcerative Colitis Endoscopic Index of Severity^11^ - and SES-CD - Simple Endoscopic Score for Crohn’s Disease^12^ - assessed by endoscopy, Bristol stool score, and the clinical activity scores HBI - Harvey-Bradshaw Index^13^ and PMS - Partial Mayo Score^14^) showed significant positive correlations (**Extended Data Fig. 1c-e**), highlighting their interconnectedness in capturing the severity and progression of IBD.

### Molecular disease activity scoring to enhance IBD assessment

The assessment of disease severity in IBD is crucial for selecting appropriate treatment regimens and adequately assessing response to initiated therapies. However, there is no universally defined and validated standard for measuring disease activity. Although existing scores demonstrate significant positive correlations with underlying severity of disease (**Extended Data Fig. 1c-e**), a definitive measure capable of identifying disease activity, including subclinical inflammation that may persist undetected at the molecular level, has yet to be established. Argmann et al.^15^ recently introduced biopsy- and blood-based molecular signatures—the biopsy molecular inflammation score (bMIS) and the circulating molecular inflammation score (cirMIS)—derived from RNA-seq data to evaluate disease severity. Following their approach, we calculated biopsy inflammatory scores for our collected samples, which effectively distinguished inflamed IBD from non-inflamed IBD and non-IBD control groups (**Extended Data Fig. 2a**). However, measuring a panel of over 100 genes, as done in the cirMIS, is impractical for routine clinical use. To address this, we developed the IBD Inflammatory Protein Severity Signature (IBD-IPSS), a more straightforward approach based on the quantification of serum proteins derived from patients’ blood. First, we performed principal component analysis for detecting potential confounding factors (**Extended Data Fig. 2b**). Subsequently, we employed the methodology outlined by Argmann et al.^15^, to conduct a differential protein abundance analysis comparing samples from actively inflamed and non-inflamed patients (**Fig. 2a**). For each of the three subtypes (IBD, UC and CD), significantly upregulated proteins were identified and incorporated into distinct inflammatory protein severity signatures: IBD-IPSS (42 proteins), UC-IPSS (32 proteins), and CD-IPSS (25 proteins), with 17 proteins shared across all conditions (**Fig. 2b**, **Extended Data Table 2**). We then compared these protein-based signatures with the cirMIS scores and found that a single protein, namely oncostatin M (OSM)^16^, was shared among all signatures (**Extended Data Fig. 2c**). To further evaluate the IBD-IPSS, we performed an in-silico protein-protein interaction analysis, which indicated that proteins from our signature are predominantly implicated in cytokine-related pathways (**Extended Data Fig. 2d**). Additionally, a protein-protein interaction network analysis identified five major clusters, all of which have been determined to be critical processes in the pathophysiology of IBD^17–20^: neutrophil chemotaxis, interleukin-6 family signaling, interleukin-7 signaling, interleukin-18 mediated signaling pathways, and positive regulation of cellular respiration (**Fig. 2c**). Since a direct comparison with blood-derived RNA-seq scores is not possible within our cohort, we evaluated the correlation between the computed IPSS-score (**Extended Data Table 3**) and several established inflammatory outcome measures including endoscopic scores (UCEIS and SES-CD), histopathology scores (modified Riley and modified Naini Cortina score), clinical activity scores (PMS and HBI), and computed molecular inflammation scores (bMIS-UC and bMIS-CD; **Extended data Table 4**). The results, presented in **Fig. 2d**, demonstrate that the serum protein signatures exhibit the strongest correlation with endoscopic scores, with a Pearson correlation coefficient (R) of 0.75 for UC-IPSS and UCEIS and R=0.58 for CD-IPSS and SES-CD. To complete the serum protein characterization, we compared inflamed and non-inflamed IBD samples with non-IBD controls (**Extended Data Fig. 2e**), confirming that OSM levels are significantly elevated during inflammation (0.58 increase in mean NPX in inflamed IBD vs. nonIBD and 0.56 increase in mean NPX in inflamed IBD vs. non-inflamed IBD samples; adjusted p-value < 0.01). Notably, TNF and AXIN1 showed a significant increase in inflamed (1.37 and 0.52 increase in mean NPX, respectively) and non-inflamed IBD (1.17 and 0.6 increase in mean NPX, respectively) compared to non-IBD controls, suggesting that these markers may serve as effective biomarkers for IBD, irrespective of the disease activity status, whether it is active or in remission.

**Fig. 2.**
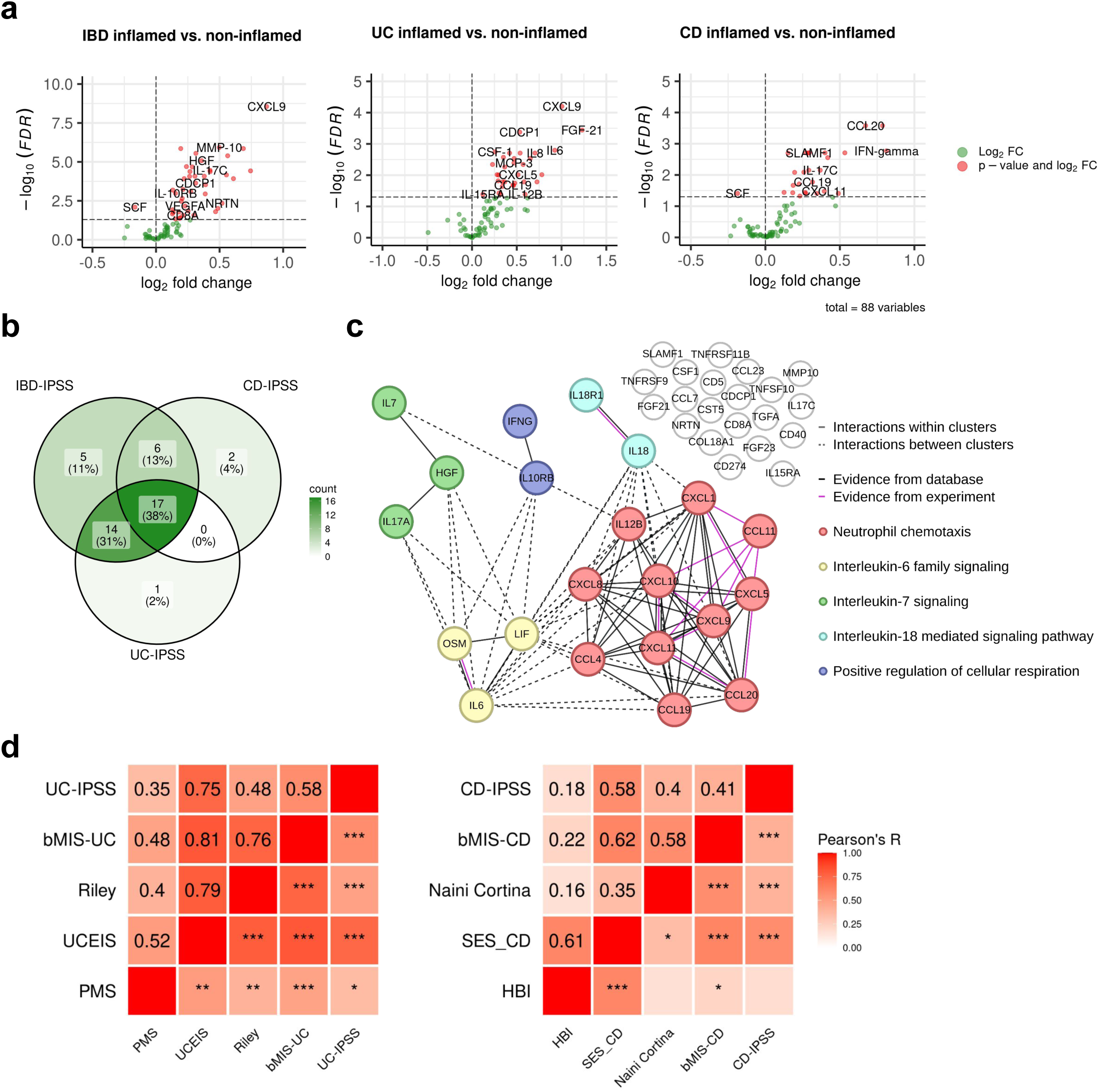
Inflammatory protein severity signature (IPSS). **a,** Volcano plots of differentially abundant serum proteins in IBD-inflamed vs. non-inflamed, UC inflamed vs. non-inflamed and CD inflamed vs. non-inflamed samples assessed by Welch t-test with an adjusted p-value <0.1. **b,** Overlap of proteins in the different inflammatory protein severity signatures. **c,** Protein-protein interaction network of the serum proteins of the IBD-IPSS. **d,** Pearson correlation of the inflammatory protein severity signatures with biopsy molecular inflammation scores (bMIS-UC and bMIS-CD) derived from gene set variation analysis from RNA-seq data, histopathology scores (normalized modified Riley score and normalized modified Naini-Cortina score), endoscopic scores (UCEIS = Ulcerative Colitis Endoscopic Index of Severity, SES-CD = Simple Endoscopic Score for Crohn’s Disease) and clinical activity scores (PMS= Partial Mayo Score, HBI=Harvey-Bradshaw Index) for UC and CD, respectively; *** p<0.001, ** p<0.01, * p< 0.05;

### Distinct immunological pathways underpin site-specific inflammatory signatures in IBD

In recent years, mounting evidence has highlighted substantial disparities between ileal CD and colonic CD across diverse intestinal layers. Colonic CD has been observed to manifest comparable disease characteristics to UC, reinforcing the notion that IBD encompasses a more intricate spectrum of disease manifestations beyond the conventional classifications of CD and UC^21,22^. The principal component analysis of RNA-seq profiles from the IBDome atlas underscored that the tissue type accounted for the largest variance (PC1=62%), followed by inflammation grade (PC2=12%). Notably, there was no clear visual separation between the overall disease entities CD and UC (**Fig. 3a**). Subsequently, we grouped the transcriptomic samples by disease entity, sampling site, and histologic disease activity (CD colon inflamed, CD ileum inflamed, and UC colon inflamed) and performed differential gene expression analyses relative to the corresponding non-IBD control groups (non-IBD colon and non-IBD ileum) (**Extended Data Fig. 3a, Extended Data Tables 5-7**). The overlapping differentially expressed genes (adjusted p-value < 0.05 and |log2FoldChange| >1) are shown in **Fig. 3b** and **Extended Data Fig. 3b**. An over-representation analysis (ORA) of these significantly upregulated genes, using the Gene Ontology - Biological Process (GO-BP) database, revealed enrichment for known immune-related pathways, including acute inflammatory response (fold enrichment=8.12), chemokine (fold enrichment=6.92), and cytokine production (fold enrichment=3.48) (**Fig. 3c**). ORA of overlapping downregulated genes did not show enrichment for any term, but the expression profiles are shown in **Extended Data Fig. 3c,** highlighting the differences in gene expression between different tissues.

**Fig. 3.**
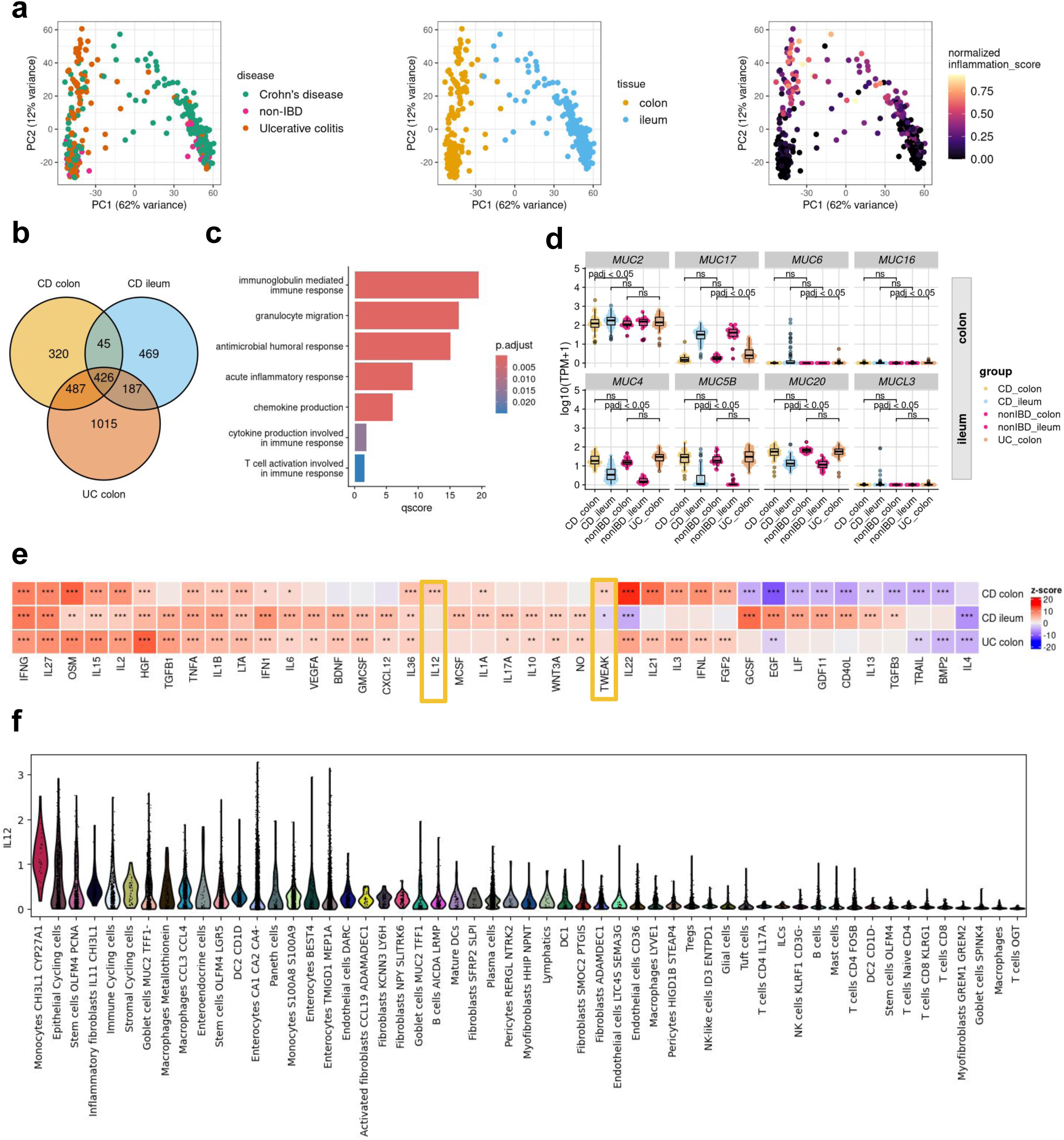
Tissue-disease-specific inflammatory gene signatures. **a,** Principal component analysis of gene expression data, colored by disease type, tissue and normalized inflammation as assessed by histopathology (normalized modified Naini Cortina score or normalized modified Riley score). **b,** Venn-diagram depicting the overlap of DE genes in the different comparisons (CD inflamed colon vs. non-IBD colon; CD inflamed ileum vs. non-IBD ileum and UC inflamed colon vs. non-IBD colon). **c,** Commonly upregulated GO-BP terms across all groups. **d,** Expression [log10(TPM+1)] of significantly upregulated MUCINs detected by DE analysis; adjusted p-values were derived from the DE analysis with DESeq2. **e,** Cytokine signaling activities in the different groups inferred with CytoSig; z-scores and p-values were derived with the CytoSig permutation test (more details in methods); * FDR < 0.1, ** FDR < 0.05 and *** FDR < 0.01. **f,** IL12 signaling activity in different cell types of inflamed CD samples (dataset from Kong et al. *Immunity* 2023).

The composition of the mucus layer varies between the colon and ileum^23^, and previous studies have shown that the structure and function of the mucosal barrier, including the mucus layer, may be significantly disrupted in IBD^24,25^. Mucins (MUCs), which are proteins expressed by epithelial cells, are key components of the mucus. Differential gene expression analysis revealed that seven mucins and one mucin-like gene were significantly upregulated (adjusted p-value < 0.05 and |log2FC| > 1): *MUC2* in the colon of CD patients, *MUC6*, *MUC16*, and *MUC17* in the colon of UC patients, and *MUC5B*, *MUC4*, *MUC20*, and *MUCL3* in the ileum of CD patients (**Fig. 3d**). *MUC6*, *MUC16*, and *MUCL3* are generally expressed at low levels and are therefore likely to be of limited relevance. In contrast, *MUC17*, a transmembrane mucin found in both the colon and small intestine, is significantly upregulated in inflamed UC colon samples compared to non-IBD controls, but no significant changes were observed in CD. Interestingly, we also observed an upregulation of *MUC4* in inflamed ileal CD samples, although *MUC4* is primarily associated with colonic membrane mucins.

To better understand the signaling pathways involved in IBD, we inferred cytokine signaling activities using CytoSig^26^. Unlike traditional approaches that rely on pathway gene expression, CytoSig infers signaling activities by focusing on the expression of genes that respond to pathway activation. The majority (n=40) of cytokine signaling pathways encoded within CytoSig (total n=43) were significantly activated or suppressed in at least one of the site-specific conditions (**Fig. 3e**). The most commonly known pathways, such as TNFA, OSM, and IFNG, show consistently high activation in all inflamed samples compared to non-IBD controls. Notably, we also identified site-specific pathway activations, including IL-22, IL-21, IL-3, interferon lambda (IFNL), and fibroblast growth factor (FGF) 2, in inflamed colon samples, regardless of disease entity. Additionally, we observed disease-subtype specific pathway dysregulation, such as the interleukin-13 pathway in CD, but not in UC (**Fig. 3e**). This aligns with the failure of anti-IL-13 antibody therapies in clinical trials in UC^27,28^. Interestingly, two signaling pathways – IL-12 and, to a lesser extent tumor necrosis factor-like weak inducer of apoptosis (TWEAK) – were significantly active in inflamed colonic CD samples. IL-12 is a key cytokine, known to initiate Th1-mediated inflammation. Examining the expression of individual genes involved in IL-12 signaling (**Extended Data Fig. 3d**), we observed a modest, but statistically significant increase in the expression of *IL12A*, *IL12B*, and *IL12RB2* in inflamed colonic samples from CD patients compared to colonic non-IBD control samples. Consistent with our findings, Dulai et al.^29^ reported in a meta-analysis of the CERTIFI and UNITI clinical trials that treatment with ustekinumab, an IL-12- and IL-23p40 antibody, was less effective in CD patients with isolated ileal-compared to colonic disease. To investigate the cell types potentially responsible for the activation of interleukin-12 signaling in colonic CD, we utilized the published single-cell dataset of Kong et al.^30^, filtering for inflamed colonic samples and inferring cytokine signaling activities at the single-cell level using CytoSig^26^ (**Fig. 3f**). The analysis revealed upregulated IL-12 signaling activity in *CHI3L1* - *CYP27A1* positive monocytes. Chitinase-3-like protein 1 (CHI3L1) is a glycoprotein associated with several diseases, including IBD^31^ and was recently identified as a neutrophil autoantigenic target in CD^32^.

### Multi-omics profiling identifies potential serum protein biomarkers for disease localization in IBD

The identification of site-specific immune signatures, mucin expression patterns, and cytokine signaling pathways in IBD underscores the complexity of its pathogenesis and highlights the need for precise, tailored therapeutic approaches. Building on these insights, the next critical step is to translate them into actionable tools for clinical application. Specifically, we sought to determine whether distinct immunological pathways driving IBD can be leveraged to identify biomarkers capable of differentiating disease subgroups. Such biomarkers could provide a basis for improved diagnosis, stratification, and personalized treatment strategies for IBD patients^33^.

Therefore, we categorized serum protein samples into three groups based on inflammatory disease localization: CD-ileum (isolated ileal disease), CD-colon, and UC-colon. We then performed a differential protein abundance analysis comparing samples from IBD patients with active inflammation against non-IBD controls (**Fig. 4a, ExtendedDataTables 8-10**). This analysis identified five proteins—TNF, IL-12B, AXIN1, OSM, and tumor necrosis factor superfamily 14 (TNFSF14)— that were commonly upregulated in all patient groups. Colon samples of both IBD entities showed the highest overlap of differentially abundant proteins (n=8: CCL20, CCL25, CXCL1, CXCL11, EN-RAGE, HGF, IL-24, and LAP TGF-beta-1), while no commonly regulated proteins were identified between ileal CD and colonic UC (**Fig. 4b, Extended Data Fig. 4a**). In ileal CD, the uniquely regulated proteins CUB domain-containing protein 1 (CDCP1), leukemia inhibitory factor receptor (LIF-R), and C-X3-C motif chemokine ligand 1 (CX3CL1) were all downregulated in patients with active inflammation compared to non-IBD controls.

**Fig. 4.**
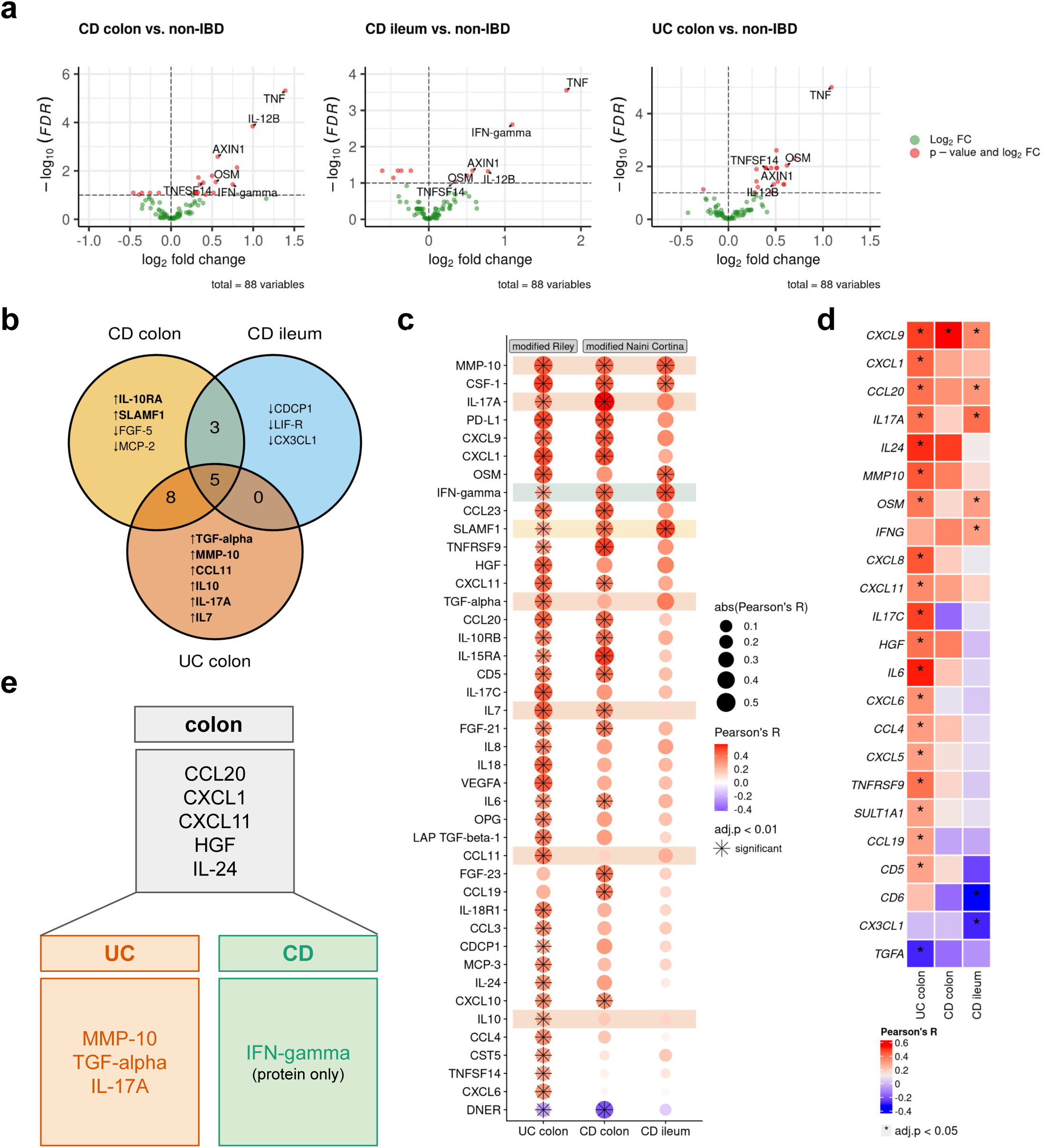
Multi-omics profiling identifies potential serum protein biomarkers for disease localization in IBD. **a,** Volcano plots displaying differentially abundant proteins in inflamed, disease-site-specific groups compared to non-IBD controls. Statistical significance was determined using Welch’s t-test with Benjamini-Hochberg correction (FDR < 0.1). **b,** Venn diagram illustrating the overlap of significantly differentially abundant proteins among CD colon, CD ileum, and UC colon, relative to non-IBD controls. **c,** Dot plot showing Pearson correlation coefficients (R) between serum protein abundance and histopathology scores (modified Riley score for UC, modified Naini Cortina score for CD) across the three subgroups. Highlighted are uniquely identified differentially abundant proteins from a and b. Significance threshold: adjusted p-value < 0.01. **d,** Heatmap of Pearson correlation coefficients between serum protein abundance and tissue gene expression in the different groups; * adjusted p-value < 0.05. **e,** Potential serum proteins associated with colonic disease, UC, and CD that significantly correlate with histopathology scores and, with the exception of IFN-gamma, also with tissue gene expression.

To explore potential associations between severity of inflammation and protein abundance, we integrated protein data with histopathology inflammatory scores of both IBD entities (modified Naini Cortina score for CD and modified Riley score for UC). In UC, all six upregulated serum proteins— Transforming Growth Factor alpha (TGF-*α*), matrix metalloproteinase-10 (MMP-10), CC-chemokine ligand 11 (CCL11), IL-10, IL-17A, and IL-7 (**Fig. 4b**) —showed significant positive correlations with the modified Riley score (**Fig. 4c**). Conversely, only one protein exhibited a significant correlation with the modified Naini Cortina score in colonic CD (SLAMF1, **Fig. 4c**). Notably, most colon-specific proteins (shared between colonic CD and UC) were also positively correlated with the histologic inflammation scores, with the exception of two proteins, CCL25 and EN-RAGE (**Extended Data Fig. 4a,b**). Among the overlapping proteins in CD, an increased abundance of IFN-gamma and decreased abundance of FGF-19 and CCL4 were observed. However, only IFN-gamma displayed a significant correlation with the severity of inflammation (**Fig. 4c**). Mucosal expression of interferon-gamma is known to be upregulated in inflamed CD^34^.

Building on these findings, we next examined the association between protein abundance in the serum and tissue gene expression. Across all samples, the strongest correlation between protein abundance and tissue gene expression was observed for CXCL9 (Pearson’s R=0.4) and the strongest inverse correlation for IL2 (Pearson’s R=-0.4) (**Extended Data Fig. 4c**). Stratification of samples by disease and site revealed several significant correlations, such as CCL20, CXCL1, CXCL11, HGF and IL-24 in colonic samples (**Extended Data Fig. 4c**) and MMP-10, IL-17A and TGF-alpha (inverse correlation) in UC (**Fig. 4e**).

Summarizing these results, we identified 5 proteins (CCL20, CXCL1, CXCL11, HGF, and IL-24) with increased abundance in colonic diseases, irrespective of the disease entity (colonic CD and UC) that significantly correlated with both, tissue gene expression and inflammatory severity. Additionally, MMP-10, IL-17A and TGF-alpha were more prominently associated with UC, while elevated serum IFN-gamma was linked to CD (**Fig. 4f**). These findings align with previous research showing higher tissue gene expression levels of *MMP10* in active UC compared to active colonic CD and controls, as well as an association with disease activity in UC^35^. Similarly, multiple studies have reported elevated HGF serum levels and mucosal gene expression in IBD, particularly in UC^36,37^.

### AI-foundation models predict accurately histologic disease activities

Histologic disease activity scoring in IBD is crucial for the assessment of treatment efficacy, prediction of disease outcomes, and for guiding clinical decision making. However, traditional scoring systems, such as the Naini Cortina score for CD and the Riley score for UC, are time-consuming, subjective and affected by inter-observer variability. In an attempt to develop a robust predictor for histologic disease activity scores, directly from pathology images of intestinal mucosal sections, we applied foundation models on images of H&E-stained tissues (**Fig. 5a**) to predict the modified Naini Cortina and modified Riley scores. Our workflow incorporates a preprocessing step where whole slide images (WSI) were tessellated into patches and color-normalized, followed by a feature extraction step leveraging four different foundation models: CHIEF^38^, UNI2^39^, Virchow2^40,41^ and H-optimus-0^42^, which is the largest open-source AI foundation model for pathology. Finally, we applied an attention-based multiple instance learning (attMIL) model to predict histologic disease activity scores (**Fig. 5a**). To evaluate the prediction performance, we used 1,212 H&E images and categorized them according to histologic disease activity scores: 699 images with the modified Naini Cortina score (514 images from Berlin and 185 from Erlangen) and 556 with the modified Riley score (472 images from Berlin and 84 from Erlangen) (**Extended Data Fig. 5a**). We performed a 5-fold cross-validation (5FCV) using the Berlin cohort (986 images in total) to train and internally validate the model.

**Fig. 5.**
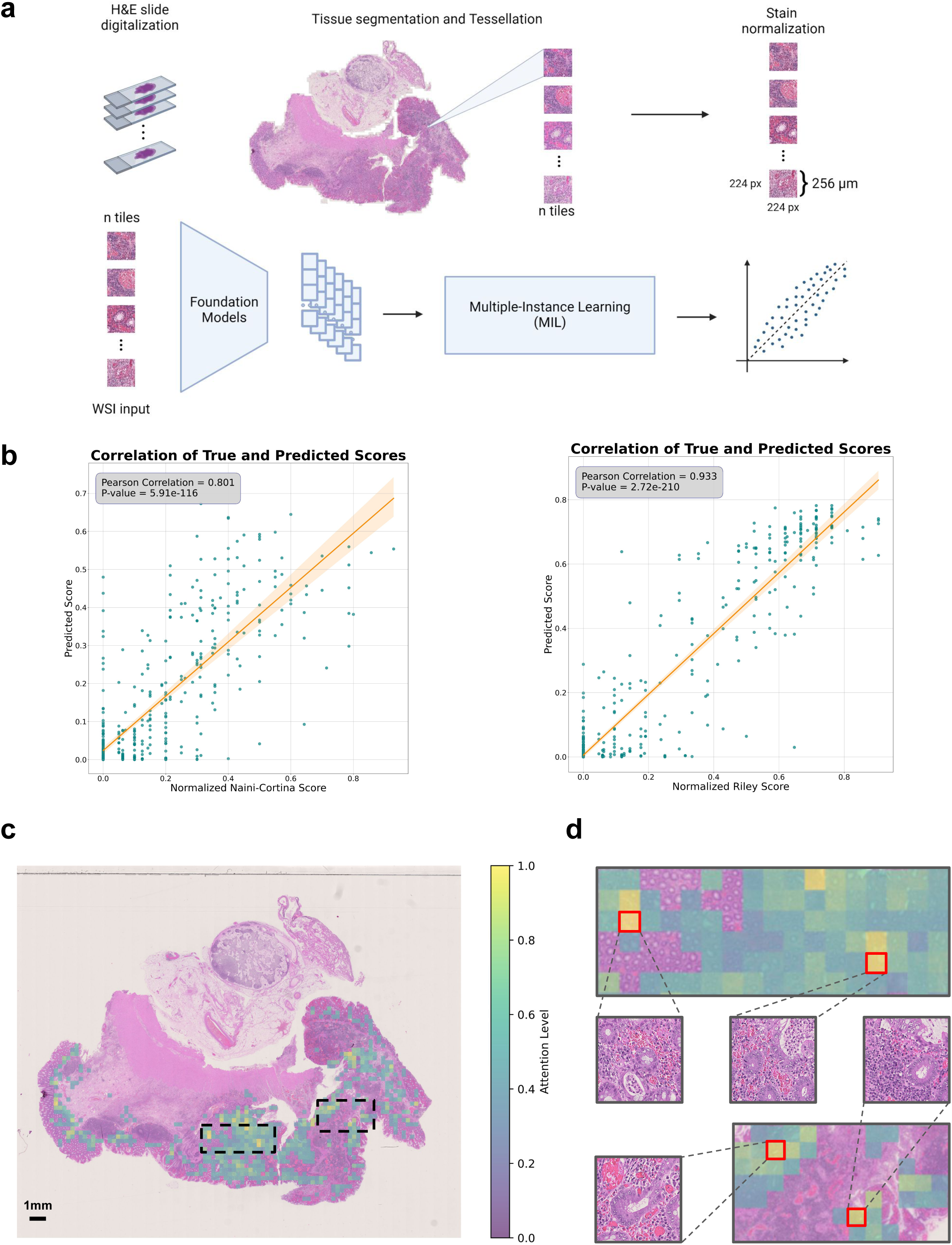
Prediction of histologic disease activity from pathology images. **a,** Overview of the image preprocessing pipeline and tile-level feature extraction, utilizing four Foundation models (CHIEF, UNI2, Virchow2 and H-optimus-0) to generate a feature matrix for each patient. An attention-based multiple instance learning (attMIL) architecture is then applied to the extracted features to predict histologic disease activity scores. **b,** Correlation plots between the original histologic disease activity scores (x-axis) and AI-predicted scores (y-axis) for both Modified Naini Cortina and Modified Riley scoring systems, based on 5-fold cross-validation on the Berlin subset using the best performing Foundation Model (UNI2 and Virchow2 respectively). **c,** Representative attention heatmap of a WSI from a UC patient with high histologic disease activity. The heatmap shows the model’s attention levels, displaying only tiles with scores above 0.4. Higher scores (yellow) mark regions that strongly influence the model’s prediction, while lower scores (green) indicate less critical regions. **d,** Zoomed-in view of the highest-attention regions highlighted in c, showing 4 of the top 10 attention tiles, outlined in red.

The performance of the different foundation models was assessed based on Pearson correlation between true and predicted scores (**Fig. 5b**). The highest performance in predicting the normalized modified Riley score was achieved by the Virchow2 model, with an R of 0.933, while the UNI2 model showed the best results for the normalized modified Naini Cortina score, reaching an R of 0.801. A comprehensive comparison of all models’ performance on the Berlin cohort across both scoring systems is provided in **Extended Data Fig. 5b**. To validate generalizability, we deployed the models to the Erlangen cohort (**Extended Data Fig. 5c**), using averaged predictions across all cross-validation folds. This approach provides a robust estimate and demonstrates strong performance achieving an R of 0.776 for the modified Naini Cortina score and an R of 0.858 for the modified Riley score.

We assessed correlations between the original (normalized modified Naini Cortina and Riley) and predicted scores with various scoring systems. While both original and predicted scores correlated strongly with bMIS in CD and UC, the predicted scores showed marginally higher correlations (CD: R=0.682 vs. 0.651; UC: R=0.799 vs. 0.790) (**Extended Data Fig. 5d,e**). Comparisons with additional scoring systems (CD-IPSS, UC-IPSS, UCEIS, SES-CD) (**Extended Data Fig. 5f**) showed that predicted scores maintained comparable or improved correlations. These findings suggest that predicted scores match or even surpass original scores, offering a viable alternative scoring method.

To understand the decision-making process of the regression model, we leveraged the attention mechanism within the attention-based multiple instance learning (attMIL) architecture. Attention heatmaps were generated to highlight the most influential regions for the model’s predictions. For detailed evaluation, we selected 10 heatmaps for each scoring system, focusing on cases with high disease activity scores and strong alignment between predicted and true scores. These heatmaps were then reviewed by expert pathologists for alignment with clinically relevant regions. In **Fig. 5c**, a UC patient’s heatmap shows the model’s attention levels. Regions with high attention (yellow) indicate strong influence on the model’s prediction, focusing primarily on peripheral areas near the mucosa and submucosa lining. These regions often display histologic signs of disease activity, such as crypt abscesses, immune cell infiltration, architectural distortion, and signs of increased epithelial regeneration, hallmarks of UC pathology. This is demonstrated by four of the top attention tiles (outlined in red) in **Fig. 5d**, which highlights areas with inflammatory cell infiltration, including lymphocytes and plasma cells as well as distorted crypts and crypt abscesses. In contrast, low-attention regions (green) are concentrated in the inner, non-inflamed, mucosal areas. Importantly, the model did not consider components of the physiologically present immune environment such as lymph follicles and lymph nodes. These results demonstrate that the model accurately identifies histologic patterns consistent with UC pathology when predicting disease activity. **Extended Data Fig. 5g** provides an additional example from a CD patient with moderate disease activity, where the model similarly focuses on pathologically relevant regions, with the top attention tile shown in the zoomed-in view.

In summary, by leveraging multiple foundation models and an interpretable attMIL framework, we show a robust and scalable solution for the prediction and assessment of histologic disease activity scores. Its high performance and generalizability can reduce inter-observer variability and enhance diagnostic accuracy in IBD.

## Discussion

We have created a comprehensive molecular, histopathologic, and clinical atlas of IBD by profiling over 1,000 patients using multi-omic and multimodal assays. Generation and integration of genomic, transcriptomic, serum proteomic, and H&E histological imaging data, coupled with standardized clinical disease characteristics annotation data, including histopathology and endoscopy scores, make IBDome a comprehensive resource for IBD. The IBDome allows the study of IBD characteristics and dissection of the phenotypic complexity in terms of molecular, cellular, and clinical features, and provides insights into the biology that could be used to improve the diagnosis and therapy of IBD. To enhance the exploitation of this resource, we are providing a publicly available, user-friendly web platform for data exploration, analysis and validation (https://ibdome.org). Beyond building this freely accessible resource for scientific research, our study provides several important insights.

First, we developed an inflammatory protein signature from serum samples that reflects the underlying intestinal inflammation and can be used to monitor disease activity of patients non-invasively. The IBD-IPSS provides a novel approach to assess disease severity, complementing existing molecular and clinical scores. Our findings demonstrate that this serum-based signature strongly correlates with established endoscopic scores, underscoring its potential as a biomarker for disease monitoring. The identification of OSM as the only overlapping protein between the IBD-IPSS and the circulating molecular inflammation score (cirMIS)^15^, suggests its central role in systemic inflammation and further supports its relevance in IBD pathophysiology^16^. While our protein-based approach offers a practical and less invasive alternative to transcriptomic intestinal tissue scoring methods such as bMIS, the clinical translation of the IBD-IPSS requires further validation.

Second, we uncovered distinct site-specific inflammatory signatures of CD and UC, emphasizing that the disease site plays a crucial role in shaping the inflammatory landscape. The observed differences between ileal and colonic CD, support the idea that IBD is more heterogeneous than the traditional CD and UC entity classification. The differential gene expression of mucins provides further insight into the tissue-specificity of IBD pathology. The selective upregulation of *MUC17* in UC colon inflammation but not in CD, and the increased expression of *MUC4* in inflamed CD ileum, suggest distinct mechanisms of barrier dysfunction in different disease subtypes. These findings highlight the need for more subtle therapeutic strategies that address the unique mucosal barrier dysfunction that occurs in different IBD subtypes. Moreover, our cytokine signaling analysis revealed key differences in inflammatory pathway activation across disease subtypes and sites. While canonical inflammatory pathways such as TNFA and OSM were consistently upregulated in all inflamed tissues, we identified site-specific and disease subtype-specific pathway activations, including IL-12 signaling in colonic CD. This is particularly relevant given the variable response to biologic therapies targeting IL-12/23, such as ustekinumab, which has been shown to be less effective in isolated ileal CD compared to colonic CD^29^.

At the serum protein level, we observed that colonic CD and UC share a substantial overlap in differentially abundant proteins, while ileal CD exhibits a more distinct inflammatory profile. The ability to differentiate IBD subtypes based on serum protein signatures offers a promising avenue for non-invasive disease monitoring and personalized treatment approaches. Specifically, the detection of MMP-10, IL-17A and TGF-alpha as UC-associated markers and IFN-gamma as a CD-associated marker may help in more accurate disease classification and targeted therapeutic strategies. Given the failure of anti-IL-13 therapies in CD^27,28^ and the ongoing investigation of anti-IFN-gamma antibodies^43,44^, our results emphasize the need to guide treatment strategies based on disease localization and immune signatures. Despite these insights, further validation in independent cohorts is necessary to confirm the diagnostic and prognostic utility of these potential biomarkers. Furthermore, the functional roles of these proteins in disease pathogenesis and their potential as therapeutic targets should also be explored further.

Third, we show that foundation models for images of H&E-stained tissue sections have superior diagnostic performance, indicating that diagnostic accuracy can be significantly improved. By leveraging several state-of-the-art foundation models (CHIEF^38^, UNI2^39^, Virchow2^40,41^, and H-optimus-0^42^) with an attention-based multiple instance learning framework, we developed a scalable and interpretable approach for predicting histologic disease activity scores with high accuracy. Our deep learning framework demonstrated high correlation between predicted and true scores, with strong generalizability across the Berlin and Erlangen cohorts. Explainability analyses showed that the model focuses on histologically relevant regions when making predictions. The attention heatmaps highlighted key pathological features closely aligning with expert pathologist assessments. Furthermore, the model’s predictions showed a strong correlation with endoscopic scoring systems such as UCEIS and SES-CD, as well as molecular scores such as bMIS and IPSS. These findings suggest that AI-based histologic scoring could reduce inter-observer variability, thereby improving objective disease monitoring in IBD and patient outcomes.

A notable limitation of our study is that although the multi-centric cohort was relatively large and complete, it lacks sufficient power for subgroup analysis. Additional studies focusing on subgroups will be necessary to increase the power. For example, stratifying patients based on disease severity (mild vs. severe) or treatment history (treatment-naïve *versus* previously treated) may provide deeper insights into disease mechanisms and therapeutic responses. We did not perform single-cell RNA sequencing or spatial single-cell analysis to further investigate cellular heterogeneity and cell-cell interactions within the tissue microenvironments of the disease localization subtypes described in this study. Spatial single-cell analysis could provide a deeper understanding of how cellular organization within tissues influences disease localization, allowing for more targeted therapeutic approaches and improved patient stratification.

In conclusion, the IBDome is a powerful resource for uncovering IBD biology and ultimately advancing precision medicine to improve patient outcomes.

## Methods

### Study centers

The IBDome study centers are located at the Department of Medicine 1, Universitätsklinikum Erlangen, and at the Department of Gastroenterology, Infectious Diseases and Rheumatology including Clinical Nutrition at the Charité – Universitätsmedizin Berlin.

### Ethics approval and consent to participate

The IBDome was approved by the institutional ethics boards in both Erlangen and Berlin (project identifiers 332-17B and EA1/200/17, respectively). The IBDome is granted permission to collect and share patient samples, clinical and molecular data. All included participants are 18 years or older and have provided informed consent before inclusion into the study.

### Data management

We distinguish between clinical databases at the study center and a centralized research database. The former was implemented by the IT departments of the study centers in accordance with data protection laws, while the latter only contains non-identifiable information that may be shared publicly according to the ethics approval. In regular intervals, data are transferred from the clinical centers to the central research database located in Innsbruck (Biocenter, Institute of Bioinformatics at the Medical University of Innsbruck).

Study participants were assigned a randomly generated pseudonym when entering the study, which was used to label specimens and samples in the research database. The data related to biomaterials are stored in pseudonymized form in the Starlims biobank management software. Access to the systems (clinical databases and Starlims) was restricted and regulated by an authorization concept.

To ensure data security, all systems are hosted in a secured environment of the university hospital IT infra-structure of Erlangen and Berlin with an information security management system (ISMS) based on guidelines from the German Federal Office for Information Security. The ISMS specifies procedures and rules within the hospital to define, manage, control, maintain, and continuously improve data security. The documented standard operating procedures for data security and data safety were followed and were checked on a regular basis. The data management fulfills all requirements of the EU General Data Protection Regulation and good scientific practice.

### Collection of clinical data

A standardized and unified medical questionnaire was designed and implemented as part of the clinical information systems of both study centers. The questionnaire consists of two parts: (1) basic data, which is entered at the initial visit, including birth year, sex, diagnosis, and pre-existing conditions, and (2) time course longitudinally collected data, which the attending doctor enters at each visit, including body weight, disease activity scores, and ongoing medication. Clinical disease activity is recorded as Partial Mayo Score (UC)^45^ and Harvey-Bradshaw Index (CD)^46^, respectively. Several consistency checks ensure data integrity during data entry.

### Biomaterial collection, processing and storage

The following specimen are collected from patients in the study

● whole blood, collected in heparinized tubes (Vacuette® Greiner Bio-One plasma tube with heparin, Thermo Fisher Scientific) for peripheral blood mononuclear cell isolation as well as K3EDTA tubes (Vacuette® Greiner Bio-One, Thermo Fisher Scientific) for DNA isolation.
● Serum, collected in (Vacuette® Greiner Bio-One Z Serum Sep Clot Activator tubes, Thermo Fisher Scientific).
● Mucosal biopsies collected during endoscopy or after surgery from surgical specimen, stored in test tubes containing RNA protect reagent (RNAprotect Tissue Reagent, Qiagen) for RNA isolation and neutral buffered, 10 % formalin solution (Sigma-Aldrich) for histopathology.
● Surgical resections, including ileocecal resection, hemicolectomy, colectomy, and normal tissue during cancer surgery, where we collected the unaffected tissue at the resection margin for IBDome.
● Stool samples, by providing patients with a stool sample tube containing RNA protect reagent (RNAprotect Tissue Reagent, Qiagen) and a questionnaire to sample stool 3-5 days after endoscopy or surgery.

In brief, samples were processed as follows. Peripheral blood mononuclear leukocytes (PBMC) are isolated from whole blood employing the SepMate™-50 (IVD) tube for density gradient centrifugation (StemCell Technologies). PBMCs are stimulated with PMA/Ionomycin and LPS or left unstimulated for 4 hours. Naїve PBMC (directly after isolation), stimulated PBMC and unstimulated PBMC (with or without brefeldin A) are fixed in Proteomic Stabilizer PROT1 (SMART TUBE Inc.) and stored at −80°C for CyTOF analysis. The supernatants of LPS-stimulated PBMC are stored at −80°C for cytokine analysis. Whole blood from EDTA tubes is stored in 1 mL aliquots at −80°C for DNA isolation. Serum is stored in 1 mL aliquots at −80°C for proteomics (Olink). After incubation of biopsies in RNA protect reagent (RNAprotect Tissue Reagent, Qiagen) overnight at 4°C, biopsies are stored individually at −80°C until RNA isolation. Formalin-fixed biopsies or resected tissue is processed by and stored at iPATH.Berlin, the core unit of Charité-Universitätsmedizin Berlin for histopathology. Stool samples in RNA protect reagent (RNAprotect Tissue Reagent, Qiagen) are stored in pea-sized aliquots or 1 mL aliquots when liquid at −80°C until analysis.

### Histopathological assessment

Formalin-fixed tissues were embedded overnight and paraffin blocks were prepared. Paraffin sections (1-2 µm) were dewaxed and hydrated in a descending alcohol series. Sections were stained with hematoxylin (Merck) and eosin (Sigma-Aldrich). Sections were dehydrated in an ascending alcohol series with xylene (Carl Roth) as intermediate and coverslipped with corbit balsam (Hecht). Histomorphology of the ileum and colon was evaluated according to modified scores based on Naini and Cortina^9^ for CD and Riley^10^ for UC. The main modification of both scores include the evaluation of resected tissue with scores for submucosal and transmural inflammation, fissures and increased lymphatic follicles. Minor modifications to the Nini and Cortina scoring system add villous atrophy and fibrosis. Also, for the Riley scoring scheme, the modifications include the scores for resected tissue as well as the scoring for ileal involvement (evaluation of infiltration with acute and chronic inflammatory cells, architectural distortion and epithelial integrity).

### Endoscopic assessment

Patients who underwent endoscopy were scored according to the Ulcerative Colitis Endoscopic Index of Severity (UCEIS) ^47^ for UC and Simple Endoscopic Score for Crohn’s Disease (SES-CD) ^12^, for CD respectively. The scoring was done based on the established criteria of both scores by experienced endoscopists at both participating centers. The endoscopists were blinded to the individual molecular date of the investigated patients.

### Stool score assessment

Stool samples were taken by the patients and shipped in RNAprotect reagent accompanied by a questionnaire. In order to classify various types of feces the Bristol stool chart was used^48^.

### Whole exome sequencing library preparation and sequencing

Total DNA was isolated from whole blood using the DNeasy Blood&Tissue Kit according to the manufacturer’s protocol (Qiagen). The concentration was measured using NanoDrop One/One (Thermo Fisher Scientific). The DNA was shipped on dry ice to the NGS Competence Center Tübingen for sequencing.

### RNA-seq library preparation and sequencing

Biopsies collected during endoscopy or from resected tissue by using a single-use biopsy forceps (Olympus) were incubated in RNA protect reagent (RNAprotect Tissue Reagent, Qiagen) and stored at −80°C. For RNA isolation, biopsies were thawed on ice and homogenized in RLT buffer (Qiagen) employing the TissueLyser LT (Qiagen). RNA was isolated, cleaned and concentrated using the RNeasy kit (Qiagen) and RNA Clean & Concentrator kit (Zymo Research). The concentration was measured at NanoDrop One/One (Thermo Fisher Scientific) and quality (RNA integrity number, RIN) at TapeStation (Agilent). RNA was shipped on dry ice to the NGS Competence Center Tübingen for sequencing.

### Serum protein assessment

An serum sample aliquot was thawed on ice for one hour and centrifuged at 3,000 rpm for one minute at 4°C. Resistand PCR-clean 96-well full skirted PCR plates (ThermoFisher Scientific, catalog number AB0800) were used with 80 µL of serum per well and sealed with adhesive tape (MicroAmp seal; ThermoFisher Scientific, catalog number 4306311). The pipetting scheme for all plates was randomized by the BIH Core Unit Proteomics. Samples were shipped on dry ice to the BIH Core Unit Proteomics, Charité, Berlin for measurements with the Olink® Target 96 Inflammation panel.

### Whole exome sequencing analysis

Germline mutations were called using a custom-built nextflow pipeline. Briefly: Whole exome sequencing raw reads were cleaned from residual adapter sequences and low-quality sequences using fastp v0.12.4^49^. The reads were then aligned to the reference genome (hg38) using BWA v0.7.17^50^. Duplicate reads were marked with sambamba v0.8.0^51^. Base-call quality score recalibration was performed with GATK4 v4.2.3^52^. Germline variants are called using the haplotypecaller program from GATK4 and Strelka2 v2.9.10^53^. Variants that were called from both algorithms were used as high-confidence variants and annotated using the Ensembl variant effect prediction (VEP v104.3) tool^54^.

To investigate *NOD2*, all mutations were filtered to retain only coding variants associated with protein-coding transcripts. Exon regions were extracted from the Gencode v33 primary assembly annotation GTF file using the R-package GenomicFeatures (v.1.56.0). A transcript database (TxDb) was created with the *makeTxDbFromGFF* function. Transcript names were retrieved using the *transcripts* function and filtered to match *NOD2* transcript IDs present in our dataset. The distribution of *NOD2* mutations was visualized using the trackViewer R-package (v.1.40.0). A lollipop plot was generated, highlighting the most frequent mutations in red.

### Transcriptomics analysis

RNA-sequencing samples from four different batches were processed with the nf-core RNA-seq pipeline version 3.4^55^. In brief, sequencing reads were aligned to the hg38/GRCh38 reference genome with Gencode v33 annotations using STAR v2.7.7a^56^. Read counts and transcripts per million (TPM) were quantified using Salmon^57^.

Differential expression analysis was performed in R v.4.4.1 with DESeq2 (v.1.44.0) using raw counts and the covariate formula ∼ *group + batch + sex + scaled age*. For comparisons between IBD inflamed and non-IBD samples *tissue_coarse* was added as an additional covariate to account for the different tissues involved. Genes were considered differentially expressed if they met an adjusted p-value threshold of < 0.05 and a |log2FoldChange| threshold of >1. For visualization of the results we used the EnhancedVolcano (v.1.22.0), ggplot2 (v.3.5.1), ComplexHeatmap (v.2.20.0), and ggvenn (v.0.1.10) R-packages.

Cytokine signaling activities for bulk gene expression data were inferred using CytoSig^26^ in Python v.3.8.20, leveraging the cytosig.v0.1 implementation available on GitHub (https://github.com/data2intelligence/CytoSig). TPM values were log-transformed as log_2_(TPM + 1) prior to analysis and used as input. CytoSig calculates the z-score by dividing the regression coefficient by the standard error. The p-values are obtained using a permutation test when the random count is > 0 or using a Student’s t-test if the random count is 0.

For cytokine signaling activities at the single-cell level we used the processed dataset from Kong et al.^30^ accessible through the Broad Single Cell Portal under accession number SCP1884. To infer cytokine signaling activities, we applied weighted means (using the *run_wmean* function implemented in the decoupler-py package^58^) with the CytoSig database retrieved from OmniPath^59^.

Biopsy and circulating molecular inflammation signatures were obtained from Argmann et al.^15^. To calculate the biopsy molecular inflammation scores (bMIS) for our samples, we applied gene-set variation analysis (GSVA)^60^ using the GSVA R-package (v.1.52.3).

### Serum protein analysis

Data tables containing normalized protein expression (NPX) values, Olink Proteomics’ arbitrary unit on log2 scale, were loaded into R v.4.4.1 and further processed with the OlinkAnalyze (v.4.0.1) R-package. Differential protein analysis was conducted using the *olink_ttest* function. Only proteins detected in at least 90% of the measured samples were included in the analysis. Statistical differences were assessed using the Welch two-sample t-test with Benjamini-Hochberg correction applied to adjust for multiple testing. Proteins were considered differentially abundant if they met a FDR threshold of < 0.05. Results were visualized using the EnhancedVolcano (v.1.22.0) R-package. Intersections were retrieved and plotted with the ggVennDiagram (v.1.5.2) or the UpSetR (v.1.4.0) R-package.

We developed an IBD Inflammatory Protein Severity Signature (IBD-IPSS) using a method consistent with the approach outlined by Argmann et al.^15^. In brief, differential protein abundance between inflamed and non-inflamed IBD samples was analyzed using OlinkAnalyze as described above, identifying significantly upregulated proteins for inclusion in the IBD-IPSS. Similarly, entity-specific signatures were generated: the UC-IPSS and CD-IPSS, derived by analyzing protein abundance separately in ulcerative colitis and Crohn’s disease samples. Correlation analysis with the various inflammatory scores available within IBDome including endoscopic scores (SES-CD and UCEIS), clinical scores (HBI and PMS), histopathology scores (modified Riley and modified Naini Cortina score) and the computed bMIS scores (bMIS-CD and bMIS-UC) was conducted using Pearson correlation with pairwise complete observations.

Functional analysis and clustering of the IBD-IPSS proteins was performed using the STRING database^61^. Evidence for protein interactions was considered only from curated databases and experimentally validated interactions. Clustering was performed using MCL (Markov Cluster Algorithm)^62^ with an inflation parameter set to 3. Clusters were annotated using the default settings of the STRING database web application. This annotation process prioritized general terms or pathways that summarize multiple specific terms and pathways, derived from various databases integrated within STRING.

### Normalization of histopathology scores

To ensure comparability between different histopathology scores (modified Naini Cortina Score and modified Riley score), we normalized the scores to a 0-1 scale, considering the tissue-specific maximum score for each disease entity (CD or UC) and sampling method (biopsy or resection). The maximum scores are listed in Table 1.

**Table 1.** Maximum histopathology scores for the modified Naini Cortina and modified Riley scores categorized by tissue type and sampling method (biopsy or resection).

| tissues | sampling method | max. modified<br>Naini Cortina score | max. modified<br>Riley score |
| --- | --- | --- | --- |
| colon, rectum, caecum | resection | 20 | 21 |
|  | biopsy | 16 | 17 |
| ileum, ileocecal valve, small intestine, anastomosis, pouch | resection | 14 | 16 |
|  | biopsy | 10 | 12 |

### The IBDome research database

A relational database was designed and implemented in the Python package sqlalchemy using SQLite as database engine. Data integrity is ensured through check constraints and foreign key validation. SQLite was chosen over other database systems, because it makes the database easy to share as a single file, does not require a server, and offers good performance for a use-case without concurrent writes. Inconsistencies in clinical data were resolved manually, and implausible entries were removed. Both clinical and molecular data were processed and imported into the database in a set of Jupyter notebooks and a custom helper library written in Python. All data loading steps are integrated into a Nextflow^63^ pipeline, which allows rebuilding the database from scratch in a single command.

### Web application

The IBDome web application is implemented in R Shiny and directly interacts with the IBDome SQLite database. Dependencies are packaged in a Docker container and a docker-compose file is provided which allows executing the app locally. Plots were generated in R using the ggplot2^64^, ggpubr, plotly, and ggbeeswarm packages. For visualization of gene expression data, transcripts per million (TPM) values were log_10_(TPM+1) transformed. P-values were computed using a two-tailed Wilcoxon test on the transformed values.

### Acquisition of high-resolution H&E images

Whole slide images of H&E stained tissue sections were scanned in two batches at different centers: MUI (Innsbruck) and Charité (Berlin). WSI from the first batch were scanned at x40 magnification using a NanoZoomer S210 slide scanner (Hamamatsu), and the analysis was performed using NDP.view2 software (Hamamatsu). WSI from the second batch were scanned at x100 magnification using a Vectra3 automated quantitative pathology imaging system (Akoya Biosciences).

### Deep Learning Inflammation score prediction

H&E WSI were tessellated into patches with dimensions of 224×224 pixels, representing a 256 µm edge length. To ensure consistent color distribution across cohorts, patches from each cohort underwent color normalization using the Macenko spectral matching technique^65^, which maps images to a standardized color space. For performance comparison purposes and to ensure the robustness of our findings, we employed four distinct Foundation models—CHIEF^38^, UNI2^39^, Virchow2^40,41^ and H-optimus-0^42^—which generated feature matrices of dimensions n × 768, n × 1536, n × 2560 and n × 1536 respectively, for each patient’s pre-processed patches. Here, n is the number of (224 ×224 pixels) pre-processed image patches obtained per whole slide image. All preprocessing steps followed the STAMP protocol^66^.

These feature matrices were then processed in an attention-based multiple instance learning (attMIL) framework^67,68^ designed for weakly supervised regression. For each foundation model, a separate attMIL model was trained using 5-fold cross-validation on the Berlin cohort to predict the normalized modified Naini Cortina score and the normalized modified Riley score. The cross-validation employed score-based stratification to maintain consistent data distributions across all folds, resulting in five models trained and tested on distinct and balanced splits. To externally validate the model’s prognostic performance, all five attMIL models from the cross-validation folds were independently deployed to the Erlangen cohort to mitigate fold-specific variability. Slide-level predictions were generated by each model and then aggregated through arithmetic averaging to produce the final prognostic scores. These steps were performed using the open-source Deep Learning pipeline “marugoto”^66,69^, with the default hyperparameters (learning rate = 0.0001, weight decay = 0.01, batch size = 1).

### Explainability of the Deep Learning model

To interpret the decision-making process of the regression models, we leveraged the attention mechanism of the attMIL architecture. High-resolution attention heatmaps were created by loading the attMIL model architectures for regression into a fully convolutional equivalent^70^ with their respective weights from the training procedure. By running inference on each patient’s WSI, we extracted the attention layer associated with the score prediction and overlaid it on the WSI, highlighting the regions of focus for the model’s predictions of the scores. For visualization, we selected the Berlin cohort to observe the model performance in predicting disease activity scores. For a more detailed evaluation, we selected the top 10 attention heatmaps for each scoring system based on prediction accuracy. These heatmaps were then reviewed by an expert pathologist, who assessed the highlighted regions for correspondence with areas of known clinical relevance.

## Data and code availability

The data can be interactively explored using the IBDome Explorer (https://ibdome.org), where also the full SQLite research database and individual data tables are available for download. Raw data and complete mutation tables are not made available due to privacy concerns, but IBD-relevant SNPs as reported by de Lange et al.^71^ are included in the IBDome database. Whole slide images of the H&E stained tissue sections are available from the BioImage Archive under accession number S-BIAD1753 (doi:10.6019/S-BIAD1753). The code for reproducing the results of this study is available on GitHub: https://github.com/icbi-lab/plattner_ibdome_2025.

## Acknowledgements

This work was funded by the Deutsche Forschungsgemeinschaft (DFG, German Research Foundation) - TRR241 375876048 (Z03; B01; to ZT, AAK, RA, BS, CB) and by the Austrian Science Fund (FWF) (I3978). A.A.K was further supported by SFB1340 372486779 (TP B06). B.S. is further supported by the German Research Foundation: CRU 5023 (project number 50474582), CRC 1449-B04 and Z02 (project number 431232613); CRC 1340-B06 (project number 372486779) and project number: 418055832. JNK is supported by the German Cancer Aid (DECADE, 70115166), the German Federal Ministry of Education and Research (PEARL, 01KD2104C; CAMINO, 01EO2101; TRANSFORM LIVER, 031L0312A; TANGERINE, 01KT2302 through ERA-NET Transcan; Come2Data, 16DKZ2044A; DEEP-HCC, 031L0315A), the German Academic Exchange Service (SECAI, 57616814), the European Union’s Horizon Europe and innovation programme (ODELIA, 101057091; GENIAL, 101096312), the European Research Council (ERC; NADIR, 101114631), the National Institutes of Health (EPICO, R01 CA263318) and the National Institute for Health and Care Research (NIHR, NIHR203331) Leeds Biomedical Research Centre. The views expressed are those of the author(s) and not necessarily those of the NHS, the NIHR or the Department of Health and Social Care. This work was funded by the European Union. Views and opinions expressed are however those of the author(s) only and do not necessarily reflect those of the European Union. Neither the European Union nor the granting authority can be held responsible for them.

## Author contributions

Conceptualization: C.P., G.S., A.A.K., Z.I.C., J.N.K., J.V.P., A.H., M.F.N., C.B., B.S., and Z.T.; Analysis of WES data: D.R. and C.P.; Analysis of RNA-seq data: C.P. and G.S., Analysis of proteomics data: C.P. and R.G.; Acquisition and analysis of 16S data: S.W.; Supervision of the image analysis: A.R.M., Z.I.C., and J.N.K.; Analysis of histopathology images: S.C.; Evaluation of histopathology predictions: M.G. and S.O.; SQLite database design and implementation: G.S.; Data integration in the SQLite database: G.S. and C.P.; Implementation of the web application: G.S., R.G., C.P. and D.R., Acquisition of high-resolution images: C.M. and A.A.K.; Supervision of sample preparation for RNA-seq, WES and Olink: A.K., R.A., and A.H.; Assessment of histopathology and stool scores: A.A.K.; Writing - original draft: C.P., Writing – review & editing, all authors; Funding acquisition, C.B., B.S., A.A.K., R.A., and Z.T.;

## Other contributing authors

TRR241 IBDome Consortium: Imke Atreya^1^, Raja Atreya^1^, Petra Bacher^2,3^, Christoph Becker^1^, Christian Bojarski^4^, Nathalie Britzen-Laurent^1^, Caroline Bosch-Voskens^1^, Hyun-Dong Chang^5^, Andreas Diefenbach^6^, Claudia Günther^1^, Ahmed N. Hegazy^4^, Kai Hildner^1^, Christoph S. N. Klose^6^, Kristina Koop^1^, Susanne Krug^4^, Anja A. Kühl^4^, Moritz Leppkes^1^, Rocío López-Posadas^1^, Leif S.-H. Ludwig^7^, Clemens Neufert^1^, Markus Neurath^1^, Jay V. Patankar^1^, Magdalena Prüß^3^, Andreas Radbruch^5^, Chiara Romagnani^3^, Francesca Ronchi^6^, Ashley Sanders^4,8^, Alexander Scheffold^2^, Jörg-Dieter Schulzke^4^, Michael Schumann^4^, Sebastian Schürmann^1^, Britta Siegmund^4^, Michael Stürzl^1^, Zlatko Trajanoski^9^, Antigoni Triantafyllopoulou^5,10^, Maximilian Waldner^1^, Carl Weidinger^4^, Stefan Wirtz^1^, Sebastian Zundler^1^

^1^ Department of Medicine 1, Friedrich-Alexander University, Erlangen, Germany

^2^ Institute of Clinical Molecular Biology, Christian-Albrecht University of Kiel, Kiel, Germany.

^3^ Institute of Immunology, Christian-Albrecht University of Kiel and UKSH Schleswig-Holstein, Kiel, Germany.

^4^ Charité – Universitätsmedizin Berlin, corporate member of Freie Universität Berlin and Humboldt-Universität zu Berlin, Department of Gastroenterology, Infectious Diseases and Rheumatology, Berlin, Germany

^5^ Deutsches Rheuma-Forschungszentrum, ein Institut der Leibniz-Gemeinschaft, Berlin, Germany

^6^Charité – Universitätsmedizin Berlin, corporate member of Freie Universität Berlin and Humboldt-Universität zu Berlin, Institute of Microbiology, Infectious Diseases and Immunology

^7^ Berlin Institute für Gesundheitsforschung, Medizinische System Biologie, Charité – Universitätsmedizin Berlin

^8^ Max Delbrück Center für Molekulare Medizin, Charité – Universitätsmedizin Berlin

^9^ Biocenter, Institute of Bioinformatics, Medical University of Innsbruck, Innsbruck, Austria.

^10^ Charité – Universitätsmedizin Berlin, corporate member of Freie Universität Berlin and Humboldt-Universität zu Berlin, Department of Rheumatology and Clinical Immunology, Berlin, Germany

## Competing interests

R.A. has served as a speaker, or consultant, or received research grants from AbbVie, Abivax, AlfaSigma, AstraZeneca, Bristol-Myers Squibb, CED Service GmbH, Celltrion Healthcare, Dr Falk Pharma, Galapagos, Johnson & Johnson, Eli Lilly, Materia Prima, MSD, Pfizer, and Takeda Pharma.

J.N.K. declares consulting services for Bioptimus, France; Panakeia, UK; AstraZeneca, UK; and MultiplexDx, Slovakia. Furthermore, he holds shares in StratifAI, Germany, Synagen, Germany, Ignition Lab, Germany; has received an institutional research grant by GSK; and has received honoraria by AstraZeneca, Bayer, Daiichi Sankyo, Eisai, Janssen, Merck, MSD, BMS, Roche, Pfizer, and Fresenius.

B.S. consulted for AbbVie, Abivax, Boehringer Ingelheim, Bristol Myers Squibb, Dr. Falk Pharma, Eli Lilly, Endpoint Health, Falk, Galapagos, Gilead, Janssen, Landos, Lilly, Materia Prima, PredictImmune, Pfizer, and Takeda; received speaker fees from AbbVie, AlfaSigma, BMS, CED Service GmbH, Dr. Falk Pharma, Eli Lilly, MSD, Ferring, Galapagos, Janssen, Pfizer, and Takeda; and received grant support from Pfizer (all the money went to an institutional account at Charité).

All other authors declare no competing interests.

---

**Extended Data Fig. 1.**
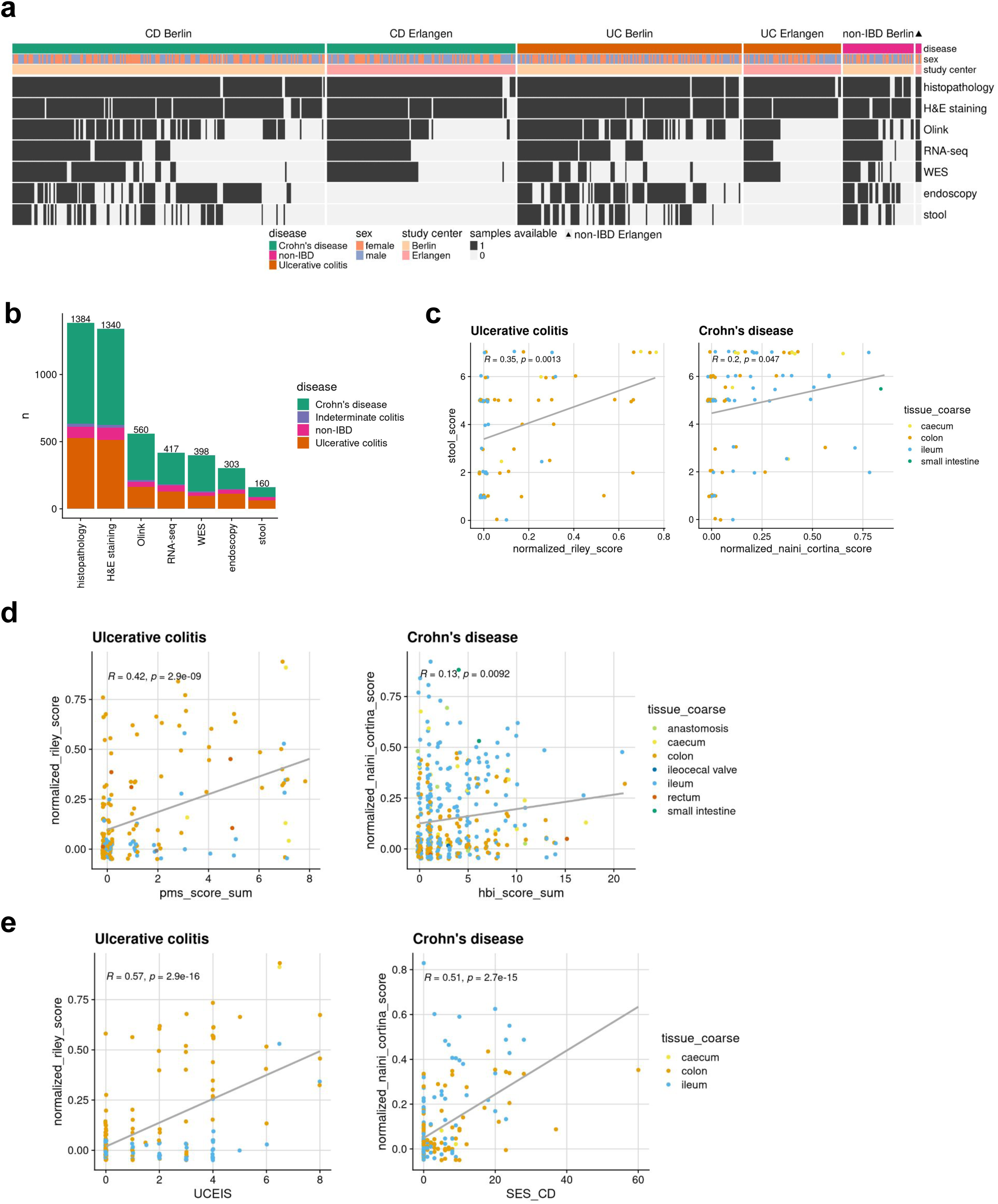
IBDome overview. **a,** Cohort distribution illustrated in a heatmap showing the available samples per patient. The heatmap is split by disease and study center and ordered according to the amount of different sample types available per patient. **b,** Number of samples per sample type; colors are representing the different diseases and number on top of the graphs are depicting the total sample numbers **c,** Correlation between histopathology (normalized Riley score and normalized Naini-Cortina score) and the Bristol stool score. **d,** Correlation between histopathology (normalized Riley score and normalized Naini-Cortina score) and clinical activity scores (PMS= Partial Mayo Score, HBI=Harvey-Bradshaw Index) for UC and CD, respectively. **e,** Correlation between endoscopic (UCEIS = Ulcerative Colitis Endoscopic Index of Severity, SES-CD = Simple Endoscopic Score for Crohn’s Disease) and histopathology scores for UC and CD, respectively.

**Extended Data Fig. 2.**
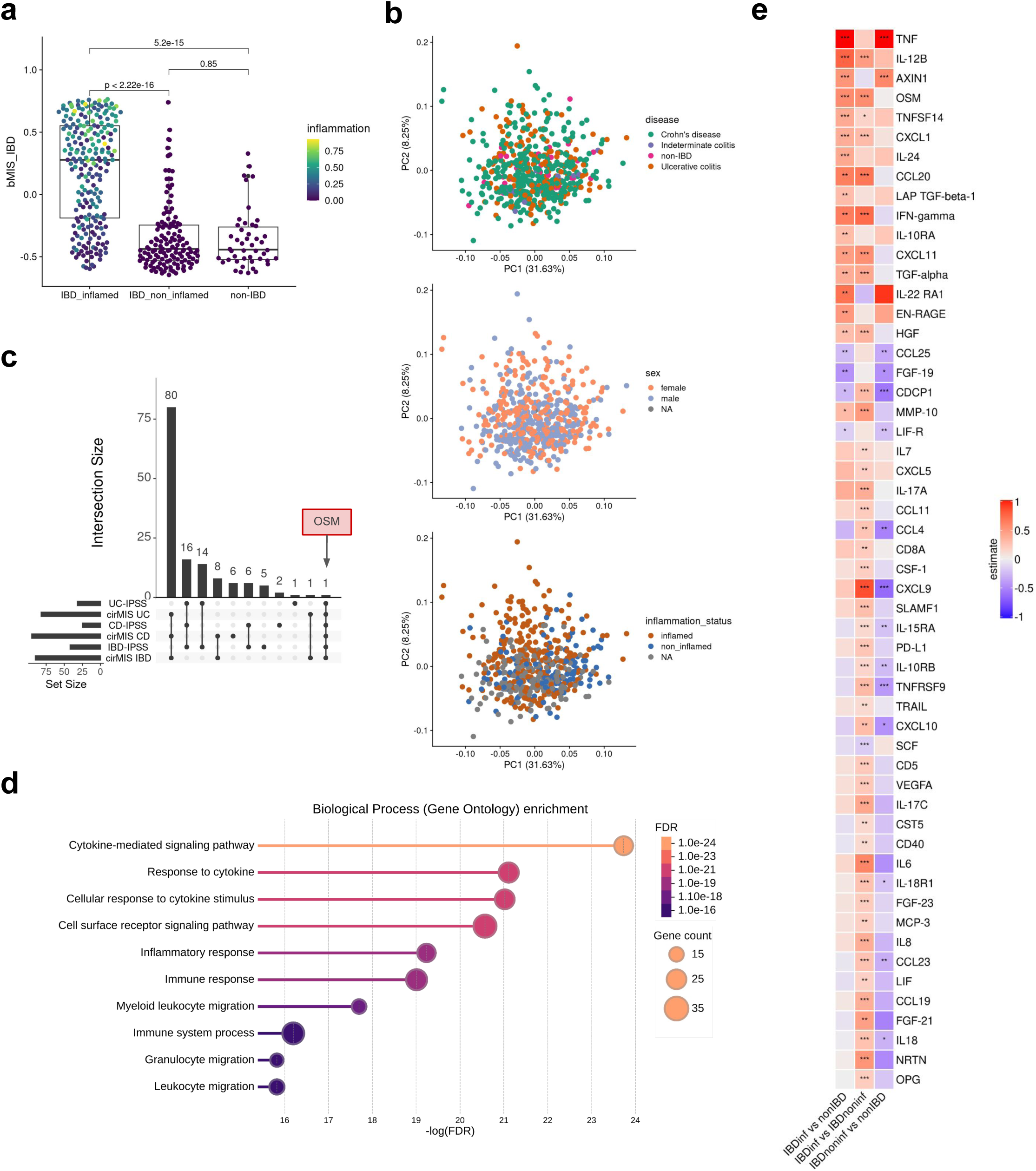
Protein abundance in the serum. **a,** Biopsy-based molecular inflammation scores (bMIS-IBD) across different groups, calculated using Gene Set Variation Analysis (GSVA); p-values were assessed using the Wilcoxon test. The color depicts the normalized inflammation score assessed by histopathology (modified Riley score for UC and modified Naini Cortina score for CD). **b,** Principal Component Analysis (PCA) of serum protein abundances, colored by disease type, sex, and inflammation status **c,** UpSet plot showing the intersection of different blood-based scores (cirMIS = circulating molecular inflammation score from RNA-seq from the blood, IPSS = inflammatory protein severity signature from serum proteins) highlighting OSM as the only protein shared across all signatures. **d,** Significantly enriched Gene Ontology - Biological Processes (GO-BP) terms (FDR < 0.05) from over representation analysis (ORA) in the IBD-IPSS. **e,** Heatmap of the differentially abundant proteins with significance levels determined by t-test: *** = adj.p<0.01, ** = adj.p < 0.05, * = adj.p < 0.1; “estimate” represent the numeric difference in mean NPX (normalized protein expression; Olink’s arbitrary unit) between groups.

**Extended Data Fig. 3.**
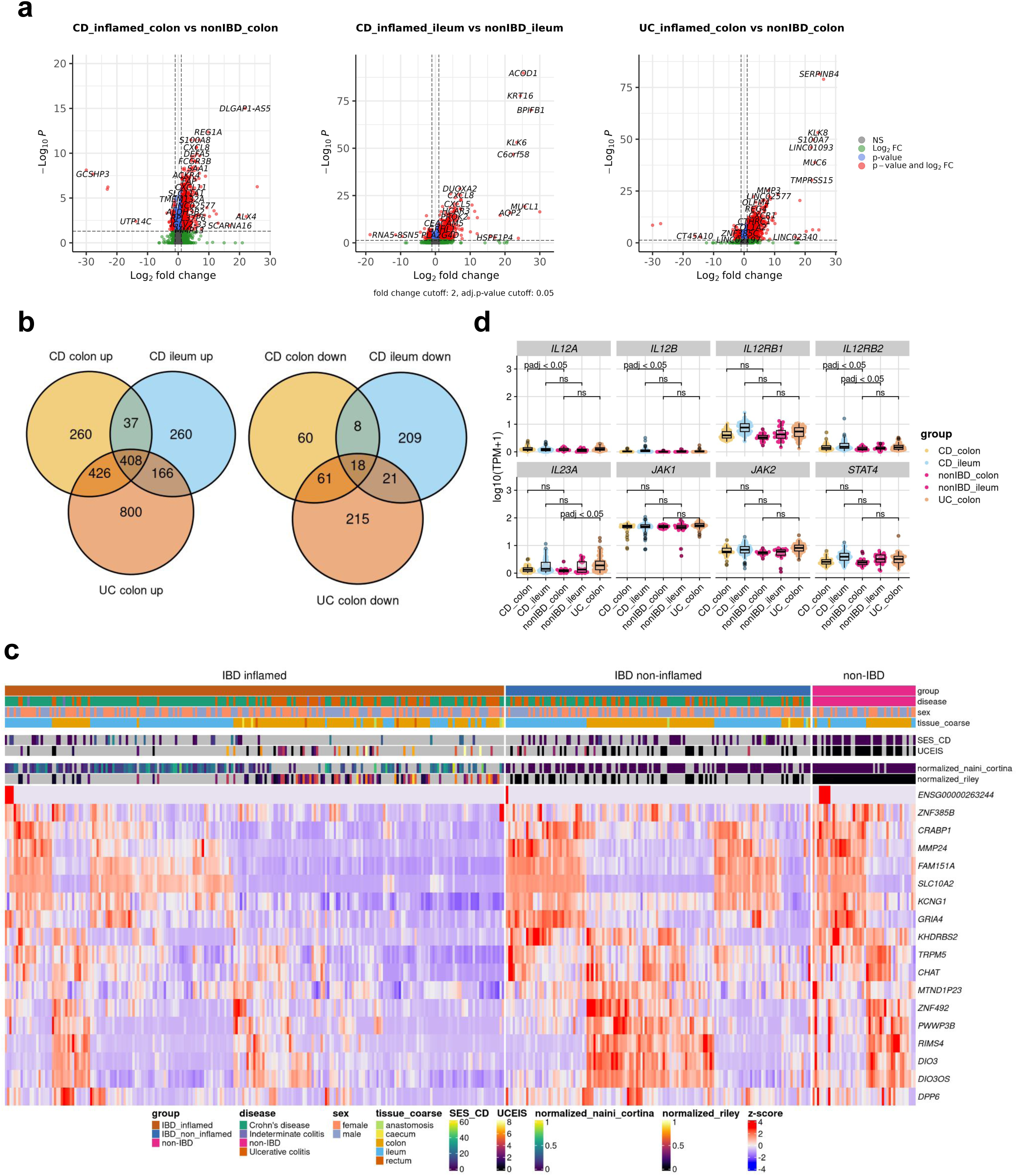
Transcriptional characterization. **a,** Volcano plots displaying the DE analysis results of inflamed CD vs. non-IBD colon samples, inflamed CD vs. non-IBD ileum samples and inflamed UC colon vs. non-IBD colon samples with thresholds: |log2FC| > 1 and adjusted p-value < 0.05. **b)** Overlap of significantly upregulated and downregulated genes in the different comparisons. **c,** Heatmap displaying the gene expression of the commonly downregulated genes (n=18) across the different groups (inflamed IBD, non-inflamed IBD and non-IBD) clustered by euclidean distance and complete linkage. **d,** Gene expression of genes involved in the IL12 signaling pathway with adjusted p-values retrieved from the differential expression analysis with DESeq2.

**Extended Data Fig. 4.**
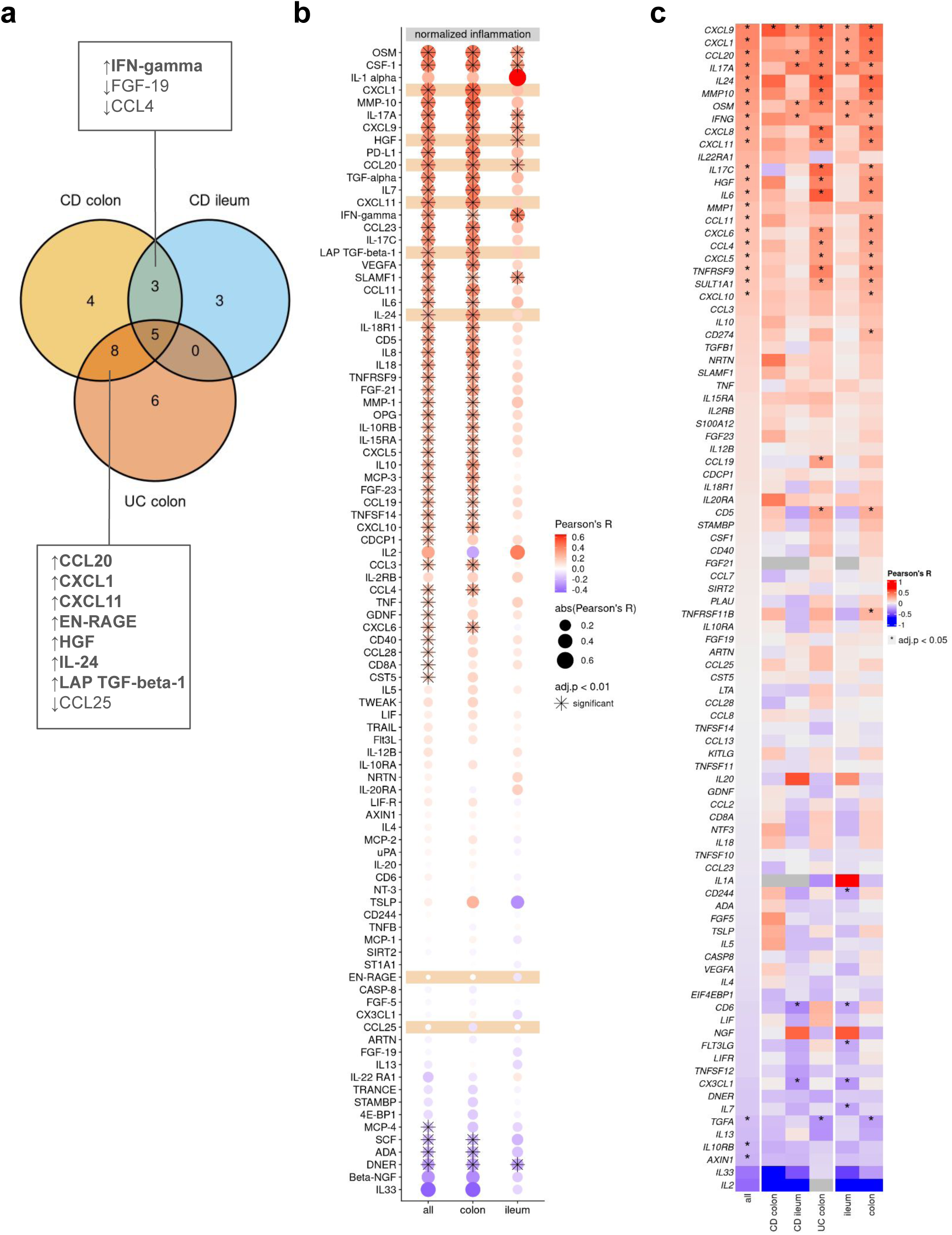
Correlation of serum proteins. **a,** Venn diagram depicting the intersections of differential abundant proteins in the different comparisons. **b,** Dot plot of Pearson correlation coefficients (R) between serum protein abundance and histopathology scores (normalized inflammation = normalized modified Riley or normalized modified Naini Cortina score) across all samples and the tissue groups; adjusted p-value cutoff: 0.01; **c,** Heatmap of Pearson correlation coefficients between serum protein abundance and tissue gene expression across all samples (n=335) and splitted by group and tissue; * adjusted p-value <0.05.

**Extended Data Fig. 5.**
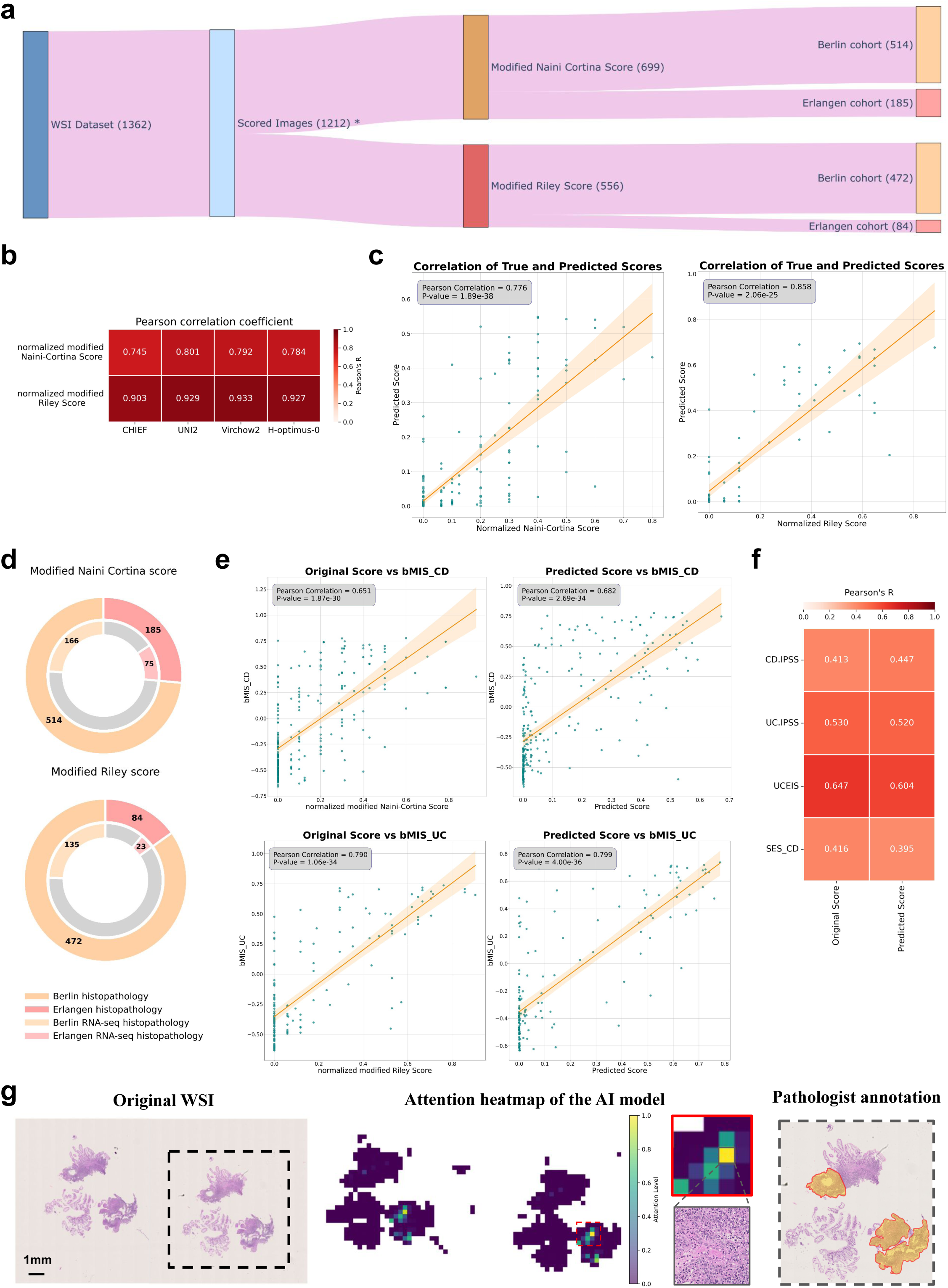
Prediction of histopathologic scores from H&E images. **a,** Distribution of all H&E images (n = 1362). After preprocessing, the number of images is reduced to 1212*, which are then categorized with two scoring systems: the modified Naini Cortina score (n = 699) and the modified Riley score (n = 556). Each scoring category is further divided in the two center subsets: Berlin (n = 514 for modified Naini Cortina, n = 472 for modified Riley) and Erlangen (n = 185 for modified Naini Cortina, n = 84 for modified Riley). *Some indeterminate colitis and non-IBD samples were scored with both scoring systems. **b,** Performance of different Foundation Models (CHIEF, UNI2, Virchow2 and H-optimus-0) on the 5 folds cross-validation regression task on the Berlin cohort, shown by Pearson correlation (R) coefficients. **c,** Correlation plots between the original histologic disease activity scores (x-axis) and AI-predicted scores (y-axis) for both modified Naini Cortina and modified Riley scoring systems. Predictions are based on the ensemble of 5 cross-validation models trained on the Berlin cohort and evaluated on the Erlangen cohort. **d,** Cohort distribution: the outer circles represent the Berlin and Erlangen histopathology cohorts, and the inner circles indicating the proportion of available RNA-seq data within each cohort. **e,** Correlation plots between the bMIS score for UC and CD with the original score (normalized Naini Cortina and Riley score) on the left and the model’s predicted scores on the right. **f,** Comparison of Pearson correlation (R) coefficients between original and predicted histologic disease activity scores against CD-IPSS, UC-IPSS, UCEIS, and SES-CD. **g,** On the left side, a representative attention heatmap of a biopsy slide image from a CD patient with high histologic disease activity. The heatmap shows the model’s attention levels. Higher scores (yellow) mark regions that strongly influence the model’s prediction, while lower scores (green) indicate less critical regions. In the middle, a zoomed-in view of the highest-attention region showing the top attention tile and on the right side the pathologist annotation highlighting the important areas to consider for disease activity scoring.

## Notes

### Summary of Updates

Section on AI-foundation models updated; Figure 5 and Extended Figure 5 updated; Supplementary tables updated; authors updated.

